# Sleeve gastrectomy protects lean mice from future obesity

**DOI:** 10.64898/2026.03.09.710623

**Authors:** Andrei Moscalu, Reji Babygirijia, Twinkle Mathew, Snehal N. Chaudhari, David Zhang, Lei Zhou, Eric G. Sheu, David A Harris

## Abstract

**Background:** Obesity and metabolic disease drive premature aging and reduced lifespan. While metabolic interventions like calorie restriction, protein restriction, and time restricted feeding have been shown to improved lifespan, they are either not effective or sustainable for most humans. Bariatric surgery is the most efficacious metabolic intervention available and is associated with increased lifespan. However, whether its longevity benefits derive solely from weight reduction or reflect surgery-specific metabolic reprogramming remains unknown.

**Methods:** We employed a lean mouse model of sleeve gastrectomy (SG) in which young, lean male C57BL/6J mice underwent SG or sham operation while maintained on low-fat chow, then were challenged with high-fat diet (HFD) in midlife. We assessed glucose metabolism, body composition, energy expenditure, hepatic histology, adipose tissue inflammation, and cecal microbiome composition.

**Results:** Despite identical weight and food intake on low-fat chow, SG mice demonstrated improved glucose tolerance and insulin sensitivity prior to HFD challenge. Upon HFD exposure, SG animals exhibited enhanced metabolic flexibility with greater capacity for fat oxidation, increased energy expenditure, attenuated weight gain, and reduced adiposity compared to sham controls. SG further reduced hepatic lipid accumulation and attenuated visceral adipose tissue inflammation, marked by decreased pro-inflammatory cytokine expression and reduced macrophage infiltration. These metabolic benefits occurred independently of caloric intake. Cecal microbiome profiling revealed surgery-specific remodeling characterized by Lactobacillus enrichment and reductions in Verrucomicrobia and Clostridia — a pattern distinct from caloric restriction and consistent with prior SG studies.

**Conclusions:** Early-life SG confers durable, weight-loss-independent protection against midlife metabolic deterioration. Gut microbiome remodeling, particularly enrichment of Lactobacillus species, represents a candidate mediating mechanism and a potential therapeutic target for aging and metabolic disease.

## 1. Introduction

Obesity and its associated metabolic comorbidities have reached pandemic proportions, driving premature aging and reduced lifespan worldwide^1,2^. Early metabolic dysfunction is a strong predictor of adult-onset disease, including late-life diabetes, cardiovascular disease, and obesity-related cancer^3–5^. Metabolic interventions like calorie restriction^6^, protein restriction^7,8^, and time restricted feeding^9^ extend lifespan however, strategies have either failed to confer meaningful metabolic benefit in humans with obesity^10^ or proven insufficiently sustainable for widespread clinical adoption.

Among available metabolic interventions, bariatric surgery has emerged as the most efficacious therapy for durable metabolic improvement. Through a single intervention in time, patients realize health benefits extending beyond weight loss with multiple studies documenting weight-loss-independent effects on host metabolism following surgery^11–15^. Sleeve gastrectomy (SG), the most commonly performed bariatric surgery, is associated with up to a decade of additional lifespan in human cohorts^16^ and leads to broad multi-organ protection, encompassing reduced cardiovascular and renal disease burden, improved pulmonary function, decreased cancer risk, and enhanced neurological health^17^. Growing evidence further indicates that surgical intervention performed at a younger age, lower body mass index, and prior to the development of advanced metabolic disease portends better long-term outcomes^18,19^. Nevertheless, a fundamental question persists: do the longevity benefits of SG derive solely from weight reduction or does surgery itself reprogram metabolic physiology in ways that confer resilience against the stressors of in midlife?

Here, we tested the hypothesis that SG performed in young, lean male C57BL/6J mice would protect against metabolic deterioration upon long-term high-fat diet (HFD) exposure in midlife. To address this, we employed our previously developed lean SG model, in which surgery performed in young mice reared on low-fat chow improves glucose metabolism, systemic and organ-specific immunity, and visceral adipose health without altering food intake, body weight, or body composition^14,20^. We demonstrate that SG confers durable, surgery-specific protection against HFD-induced metabolic deterioration, characterized by enhanced metabolic flexibility, attenuated adipose tissue inflammation, and reduced hepatic steatosis. Critically, these effects occurred in the absence of differences in caloric intake, indicating that SG operates through mechanisms distinct from caloric restriction.

We further demonstrate that SG induced remodeling of the cecal microbiome, characterized by reductions in Verrucomicrobia and Clostridia and enrichment of Lactobacillus species. In contrast, caloric restriction expands Verrucomicrobia^21,22^ underscoring the surgery-specificity of SG-associated microbiome changes. Lactobacillus species are known to favorably influence fasting glucose ^23^, body weight^24^, and hepatic steatosis^25^, and decline precipitously by midlife^26^. The SG-associated preservation of Lactobacillus abundance may therefore represent a mechanistic link between surgery, microbiome remodeling, and the observed metabolic benefits.

Collectively, these findings demonstrate that early-life SG confers durable, surgery-specific protection against midlife metabolic deterioration and identify gut microbiome remodeling, particularly Lactobacillus enrichment, as a candidate mediating mechanism. Defining how SG exerts these effects may reveal novel therapeutic targets for the treatment of aging and age-related metabolic disease.

## 2. Materials and Methods

### Animals

Lean, male C57Bl/6J mice were purchased from Jackson Laboratories (Bar Harbor, ME) at the age of 11 weeks. They were acclimated for 1 week prior to surgery. Animals were housed in climate-controlled rooms with 12-hour light and dark cycles. All animals were cared for according to guidelines set forth by the American Association for Laboratory Animal Science and all procedures were approved by the Institutional Animal Care and Use Committee at Brigham and Women’s Hospital.

### Surgical procedures and timeline

**Figure 1** shows a graphical depiction of the study design. At 12-weeks of age, mice were weight matched and either underwent SG or Sham operation as previously described^14,27^. In brief, mice were placed on a recovery gel diet (Clear H_2_0, Westbrook, ME) from 48 hours prior to surgery to 6 days post-surgery before returning to solid diet. SG was performed with a linear cutting stapler. Sham procedure consisted of short gastric vessel division and manipulation of the stomach between forceps. After recovery, mice were returned to a low-fat diet (chow; 4.5% calories from fat; Pico 5053; Lab Diet, St. Louis, MO). Weights, glucose tolerance, and body composition were tracked as below. Mice were transitioned to high fat diet (HFD; 60% calories from fat; RD12492; Research Diets; New Brunswick, NJ) on post-operative day 60. Energy expenditure was determined before, during, and after diet transition via indirect calorimetry (below). Additional metabolic phenotyping was performed during HFD exposure prior to euthanasia on POD 140, 80 days after initiation of HFD. We completed two biologic experiments with identical set up. Cohort 1 started with 5 shams and 7 SG and cohort 2 with 5 shams and 15 SG. In cohort 1 there were three SG deaths. One died on post-operative day 5 from a gastric leak and was excluded from all data. One died after oral glucose tolerance testing and one during insulin tolerance testing post-diet transition and these were excluded from all data following. In Cohort two there were 2 SG deaths. The first, on post-operative day (POD) 5 from an abscess formation and this animal was removed from all analysis. The second was again during oral glucose tolerance testing (OGTT) after diet transition and this animal was included in all data prior to death, excluding the OGTT.

**Figure 1.**
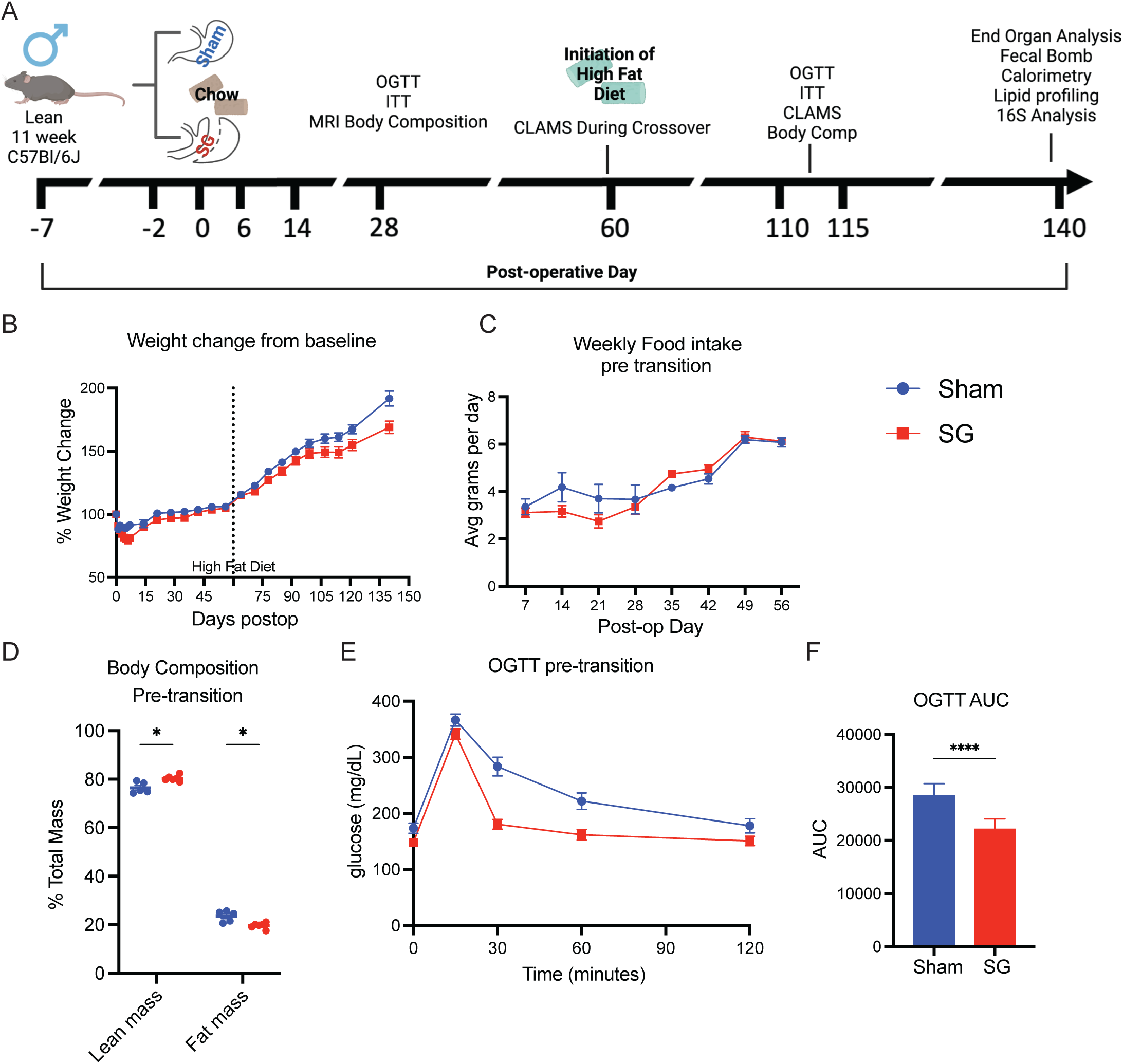
SG improves glucose tolerance in lean mice. **(A)** Experimental Design. Created in BioRender. Moscalu, A. (2024). **(B)** percent weight change from baseline across the experimental duration **(C)** Average weekly food intake pre-diet transition. **(D)** Lean and Fat mass was determined by echoMRI **(E-F)** Oral glucose tolerance and calculated AUC. (**B-F**) n from each measurement is detailed in supplemental table 1. Groups were compared using Welch’s T-test. AUC was calculated using the Trapezoid rule. Data are represented as mean +/- SEM. *p<0.05, **p<0.01; ***p<0.001; ****p<0.0001.

### Body composition and indirect calorimetry

Body weights were tracked longitudinally. Food consumption was measured weekly prior to diet switch and then daily for 1 week after diet transition. Analyses were performed at the Brigham and Women’s Hospital Metabolic Core. An EchoMRI system (Echo Medical System, Houston, Texas) was used to measure body composition immediately prior to diet transition and again prior to euthanasia. Organismal energy expenditure was measured in Comprehensive Lab Animal Monitoring Systems (CLAMS; Columbus Instruments, Columbus, OH) from POD 56 to 64 and then again from POD 94 to 101 to capture the metabolic phenotype of SG mice during initiation of and after long-term exposure to HFD. Cages were maintained at 22°C under normal 12-hour light (Fasting) and dark (Feeding) cycles. Mice were given *ad libitum* access to food and water.

### Functional Glucose Tolerance Testing

OGTT and ITT were performed after a 4 hour fast (12:00h). For OGTT, D-Glucose (2mg/g; Sigma) was delivered by oral gavage and blood glucose was measured from the tail vein a time 0 (pre-gavage), 15, 30, 60, and 120 minutes via a OneTouch Glucometer (Life Scan Inc.). For ITT, regular human insulin (0.6u/Kg; Eli Lilly) was delivered intraperitoneally. Blood glucose was measured at 0 (pre-gavage), 15, 30, 45, and 60 minutes.

### Bile acid analysis and fecal bomb calorimetry

Cecal BA analyses were performed on an Ultra-high Performance Liquid Chromatography-Mass Spectrometry instrument as reported previously ^28^. Standards were purchased form Cayman Chemicals, Sigma, or Steraloids. For each BA, we determined the standard curve and limit of detection as previously^28,29^.

### Histology

Hepatic and adipose tissue was fixed in 4% paraformaldehyde for 24 hours at 4°C, processed in 30% sucrose solution, and then embedded into Optimal Cutting Temperature compound over liquid nitrogen. Oil Red O staining was completed by the Beth Israel Deaconess Hospital Histology core facility (Boston, MA). Metabolic Associated Fatty Liver Disease (MAFLD) Activity score and Fibrosis staging were scored as outline in **Supplemental Table 2** All histologic analysis were completed by a blinded observer, Dr. Lei Zhou.

### Quantitative PCR

Tissue was snap frozen in liquid nitrogen at the time of harvest. Total RNA was extracted with Qiagen RNeasy mini kits (Germany) with an on-column DNA digestion followed by cDNA synthesis using SuperScript III First-Strand Synthesis System (Life technologies, CA). Reverse transcribed cDNAs were quantified using a SYBR green reporter. The 2^-ΔΔCt^ method was used to calculate the relative chance in gene expression. *Beta actin* was used as a housekeeping. A list of primer sequences used are found in **Supplemental Table 3**.

### Microbiome analysis

Cecal specimens were collected at euthanasia and snap frozen in liquid nitrogen. 16S sequencing and analysis were performed at Zymo Research Corporation (Irvine, Ca).

### Statistical Analysis

Two biologic replicate experiments were completed. Where appropriate, the data was combined. All values are expressed as the mean ± the standard error of the mean. Student’s unpaired Welch’s *t* tests were used to compare continuous variables. General linear modeling with lean or total mass covariates were used to compare CLAMS caging data as indicated. CLAMS data was processed in the open-source program, CALR (Available at https://CalRapp.org/) by the core facility^30^. Microbiome analysis was completed as above. All other statistical analyses were performed in Prism 10 (Graph Pad, San Diego, CA).

## 3. Results

### SG protects against glucose intolerance independent of weight loss

We have previously shown that SG improves glucose handling, insulin tolerance, and induces an organismal shift in glucose utilization independent of changes in weight ^14^. However, how these changes influence metabolic health in the face of obesogenic insult remains unclear. 11-week-old C57BL/6J mice reared on standard, low-fat chow were weight matched and underwent SG or Sham surgery followed by continued chow diet (**Figure 1A**) until post-operative day 60. As previously reported, prior to initiation of HFD, SG and Sham animals had identical weight (**Figure 1B**) and food intake (**Figure 1C**). However, MRI body composition analysis revealed that SG animals had a small but statistically significant increase in lean mass and decrease in fat mass compared to sham counterparts on post-operative day 56 (**Figure 1D**). As shown previously, despite similar anthropomorphic features, SG animals had a dramatic improvement in glucose handling in response to oral challenge (**Figure 1E, 1F**). Chow Sham and SG mice have equivalent fasting (160±33 vs 140±38, p=0.39) and peak (311±32 vs. 333 ±27, P=0.31) serum glucose levels before and during oral glucose challenge, respectively. However, chow SG mice have a more rapid descent to baseline serum glucose levels by 30 minutes (237±78 vs 117.5±31, p=0.007).

### SG protects against high fat diet induced obesity and metabolic dysfunction

Within days of exposure to an obesogenic diet (HFD; 60% calories fat), there is a clear divergence in weight gain between SG and sham animals (**Figure 1A**). HFD Sham animals have a nearly 30% increase in weight gain compared to HFD SG animals at sacrifice (POD 140 and 80 days after HFD initiation; **Figure 2A**). Further, HFD SG animals continue to have numerically greater, although non-significant, percent lean and reduced percent fat mass compared to HFD Sham mice (**Figure 2B**). These changes occur despite equivalent or, at times, higher caloric intake in SG animals (**Figure 2C**).

**Figure 2.**
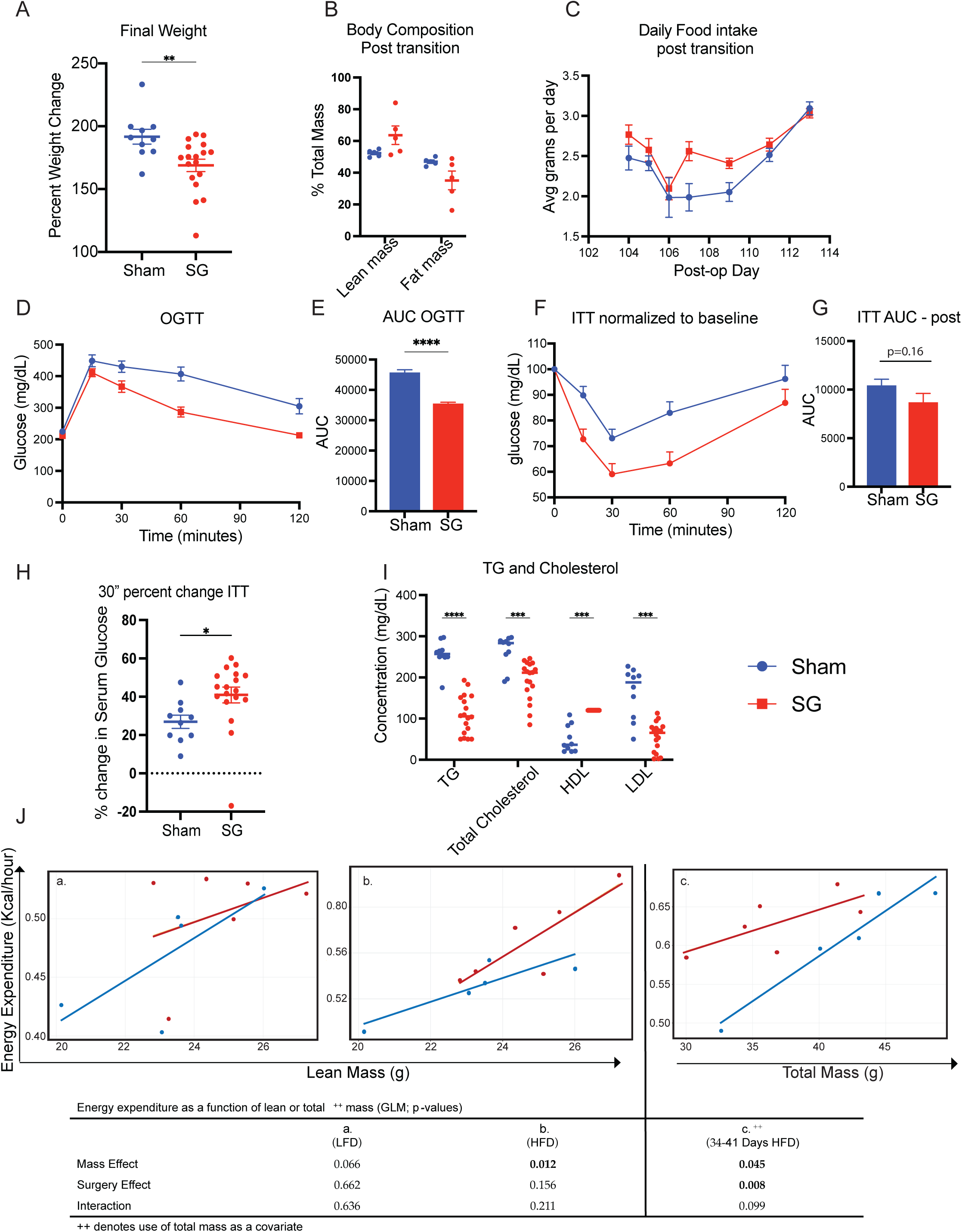
SG protects against weight gain and metabolic disease after HFD Challenge. **(A-C)** Final body weight change (A), body composition (B), and food intake (C) after diet switching. **(D-E)** Oral glucose tolerance (D) with AUC (E). **(F-H)** Insulin tolerance (F) and AUC (G) after diet transition. Glucose and insulin tolerance with corresponding AUC after diet transition. The most pronounced change in systemic glucose level occurred within the fisrt 30 minutes (H) **(I)** lipid and cholesterol profiling. **(J)** Energy expenditure before, during, and after diet transition. General linear modeling with lean or total mass covariate as indicated. (**A-I**) n from each measurement is detailed in supplemental table 1. Groups were compared using Welch’s T-test. AUC was calculated using the Trapezoid rule. Multiple comparison corrected by Holm-Sidak method. Data are represented as mean +/- SEM. *p<0.05, **p<0.01; ***p<0.001; ****p<0.0001.

Compared with HFD Sham mice, HFD fed SG mice had reduced fasting glucose (231±17 vs 187±22, p=0.007) and continued improvement in glucose tolerance (**Figure 2D, 2E**). Again, while peak serum glucose level was similar between groups, there was a more rapid decent to baseline in SG animals. SG animals were also more sensitive to intraperitoneally delivered insulin (**Figure 2F**). While there was a strong trend toward improved insulin tolerance across the entire testing period (**Figure 2G**), there was a clear, heightened response to insulin in SG mice within the first 30 minutes of testing (**Figure 2H**), suggesting heightened early glucose disposal. Lastly, HFD SG mice had improvement in serum lipid and cholesterol indices with reductions in triglyceride, total cholesterol, and LDL cholesterol as well as heightened HDL cholesterol (**Figure 2I**).

### SG mice increase energy expenditure when exposed to HFD

We placed mice in metabolic chambers during two separate times to examine how diet switching and surgery impact substrate utilization and organismal energy balance. Mice were placed in metabolic housings to capture their metabolic phenotype first during the week of transition from chow to HFD (POD 56 to 64) and again after multiple weeks of HFD exposure (POD 94 to 103).

During the first week of analysis, SG and sham mice were transitioned to HFD at POD 60. While on chow diet, there were no difference in the respiratory exchange ratio, food consumption, and locomotor or ambulatory activity (**Supp Table 4**). Additionally, there was no difference in average energy expenditure between SG and Sham animals (24 hour, fasting, feeding periods; **Supp Table 4**) or when controlling for lean mass (GLM; **Figure 2Ja**). Halfway through this week, chow diet was replaced with HFD. While there were no differences in food consumption or activity (**Supp Table 4)**, SG animals had reduced RER toward 0.7 consistent with lipid resource utilization during fasting (p=0.048), feeding (p=0.099), and over a 24-hour period (p=0.056). During the period of diet transition, SG animals did not have heightened energy expenditure (**Supp Table 4**; **Figure 2Jb**). After long term HFD exposure, the differences in RER between SG and sham animals vanished. Food intake and ambulatory activity also remained similar between groups. However, with increased HFD exposure, there was an increase in the average hourly energy expenditure of SG mice during light, dark and 24-hour periods and when controlling for body weight (**Supp Table 4**; GLM **Figure 2Jc**).

### SG protects against eWAT inflammation and hepatic steatosis

HFD Sham and SG animals were sacrificed at middle age – 8 months of age, 140 days after surgery, and 80 days after high fat diet exposure. There were no differences in the inguinal, brown, or epididymal fat pad weights between groups (**Figure 3A**). However, SG mice had reduced pro-inflammatory phenotype with a reduction in TNF-α and CCL-2 (**Figure 3B**) and a trend toward reduced crownlike structures (**Figure 3C**). We next found a 1.6-fold reduction in relative (4.8±1.2 vs 3.0±0.7%; p=0.04) liver weight in HFD SG animals (**Figure 3D**). This was accompanied by a reduction in steatosis (**Figure 3E; 3F**) without a clear impact on inflammation, ballooning, or fibrosis, which are hallmarks of Metabolic Associated Steatotic Liver Disease (MASLD). The latter features of MASLD are not typically observed in this model without additional dietary or genetic modifications^31^.

**Figure 3.**
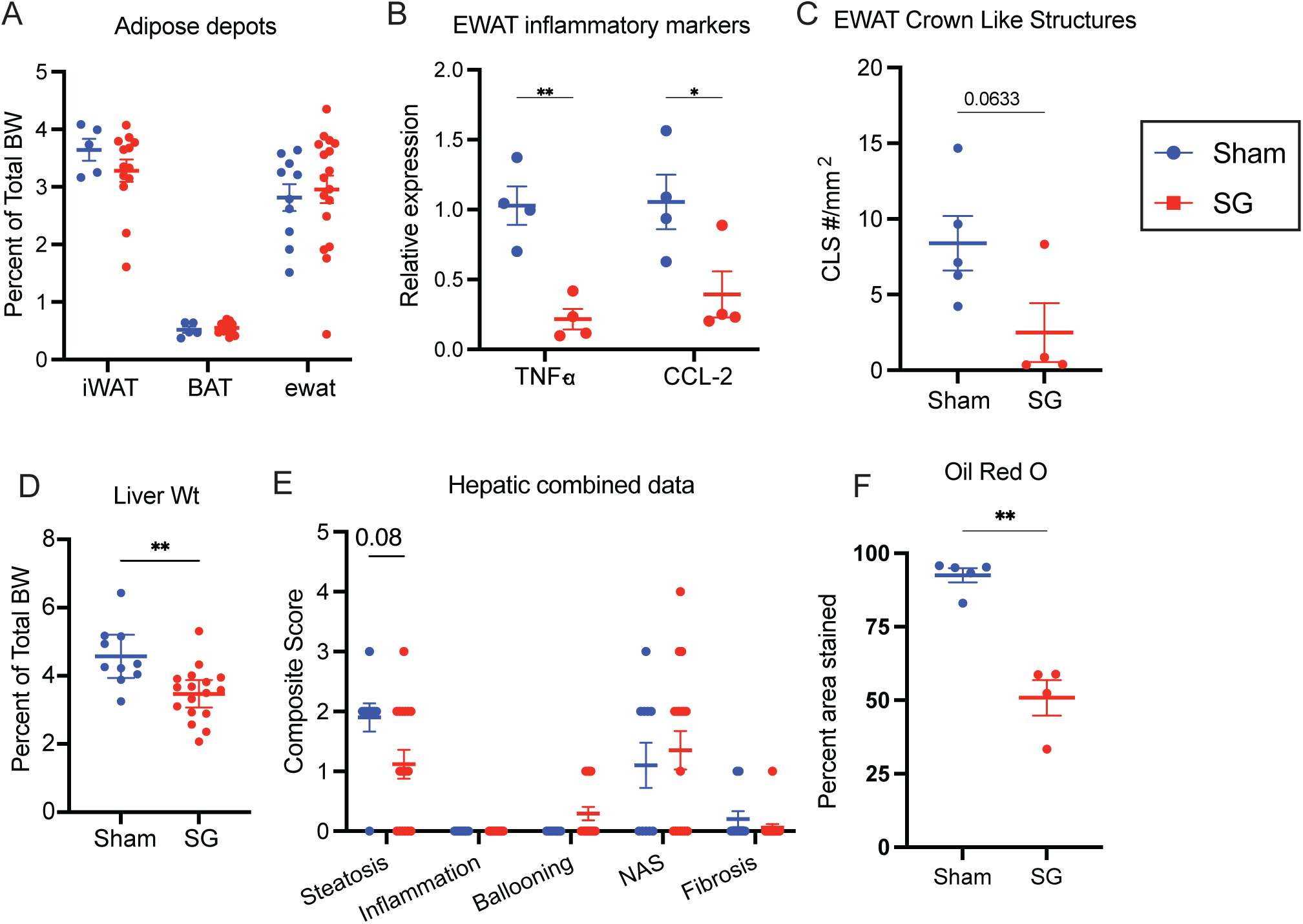
SG reduces visceral adipose inflammation and hepatic steatosis upon HFD challenge. **(A)** inguinal, brown, and epididymal adipose tissue weight **(B-C)** Epididymal inflammatory markers (B) and crown like structure density (C). **(D-F)** Total liver weight (D), NAS Scoring (E), and percent adiposity (F) (**A-F**) n from each measurement is detailed in supplemental table 1. Groups were compared using Welch’s T-test. Multiple comparison corrected by Holm-Sidak method. Data are represented as mean +/- SEM. *p<0.05, **p<0.01; ***p<0.001; ****p<0.0001.

### SG does not induce malabsorption

Given the reduced weight over time without changes in food intake, we next assessed whether SG mice have reduced nutrient absorption when exposed to HFD. To our surprise, SG mice had increased nutrient absorption with an overall reduction in the calorie per gram of stool by bomb calorimetry (**Figure 4A**). Further, there was no difference in fecal fat content (**Figure 4B**) despite there being a trend toward reduced total cecal bile acid content (**Figure 4C**). There were no statistically significant differences in individual primary or secondary bile acid species (**Figure 4D**). Thus, the reduced weight in SG mice appears to be the result of heightened energy expenditure as opposed to malabsorption.

**Figure 4.**
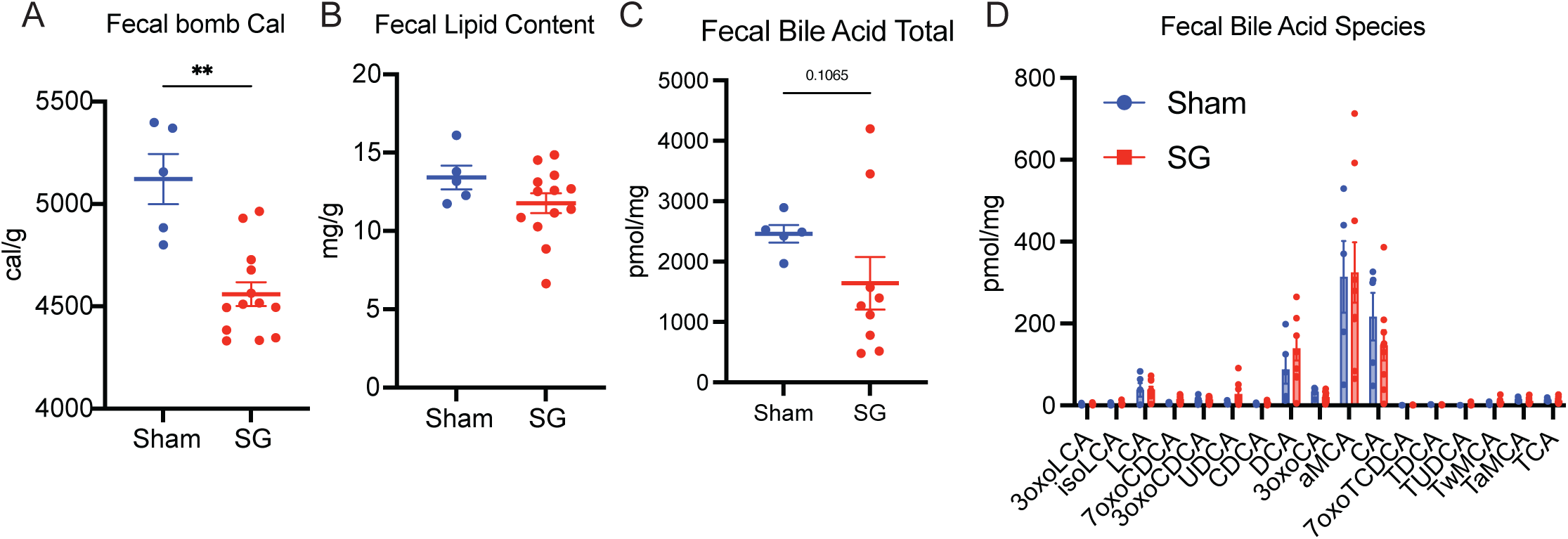
SG does not cause malabsorption. **(A)** Fecal bomb calorimetry **(B)** Fecal lipid content **(C, D)** Total cecal bile acid content (C) and individual bile acid species (D) (**A-D**) n from each measurement is detailed in supplemental table 1. Groups were compared using Welch’s T-test. Multiple comparison corrected by Holm-Sidak method. Data are represented as mean +/- SEM. *p<0.05, **p<0.01; ***p<0.001; ****p<0.0001.

### Lean SG mice exposed to HFD have a unique microbiome

The microbiome is a major contributor to host metabolic disease, and bariatric surgery induces a major shift in the intestinal microbial community that durably influences host metabolism and organ function^32–38^. Using 16s metagenomic sequencing, we found that SG mice exposed to HFD had increased absolute cecal microbial abundance compared to HFD sham animals (**Figure 5A**). There were no differences in Shannon or Chao index (**Figure 5B, 5C**) but like prior studies, we found that SG induced an increase in beta diversity of the cecal microbiota (**Figure 5D**).

**Figure 5.**
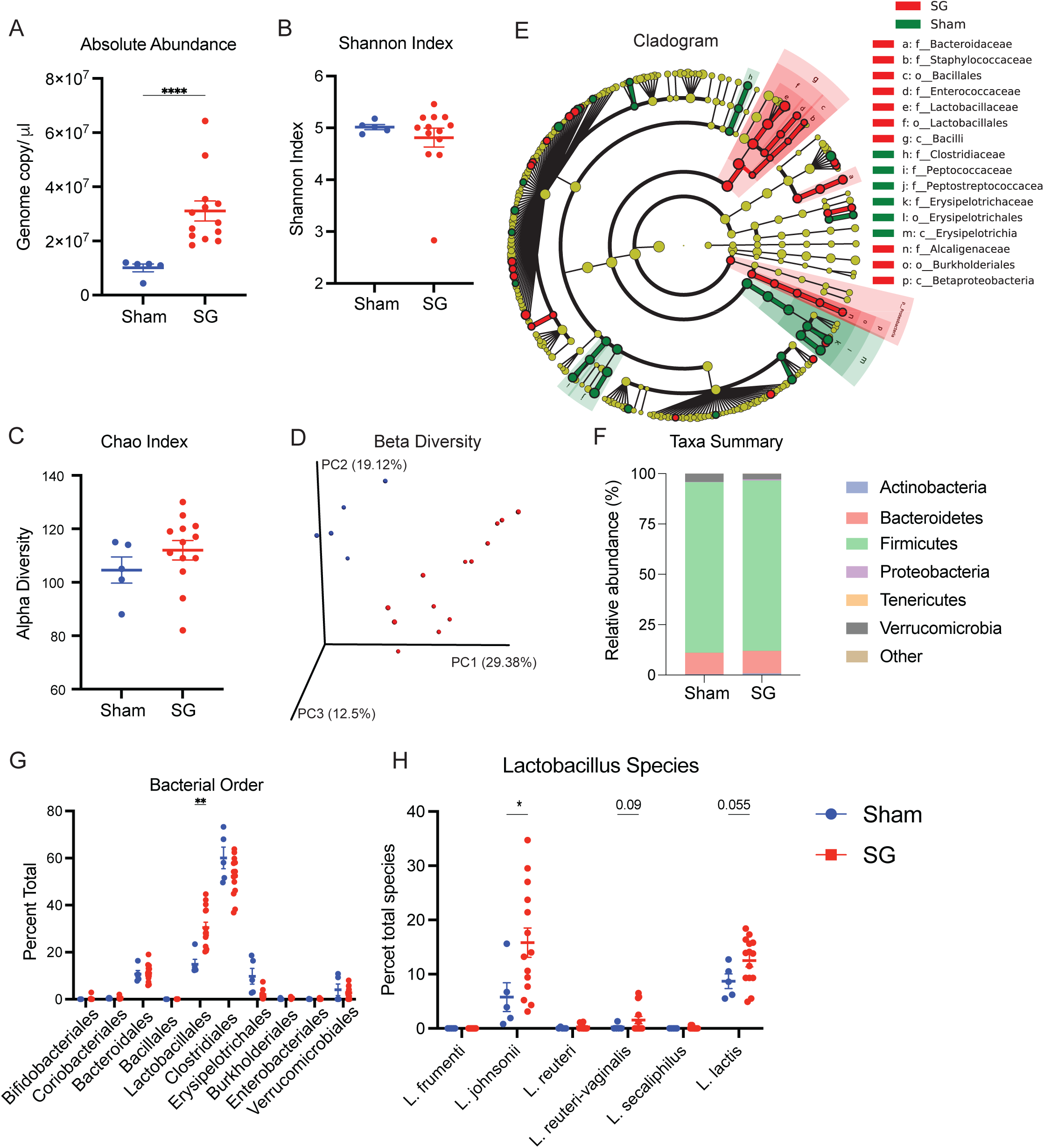
SG influences the cecal microbial composition and increases lactobacillus richness. **(A)** Absolute abundance **(B)** Shannon index of biodiversity **(C)** Chao index of species richness (D) Beta diversity (E-G) Cladogram (E), taxa summary (F), and bacterial order (G) differences between SG and sham mice after diet switching. (I) Lactobacillus species abundance. (**A-H**) n from each measurement is detailed in supplemental table 1. Groups were compared using Welch’s T-test. Multiple comparison corrected by Holm-Sidak method. Cladogram created through Zymo Research. Data are represented as mean +/- SEM. *p<0.05, **p<0.01; ***p<0.001; ****p<0.0001.

Lean SG animals exposed to HFD had a distinct cecal microbial community compared to lean sham animals exposed to HFD (**Figure 5E**). Like we have shown previously^39^, we found a numerical increase in Firmicutes, Bacteroidetes, and Proteobacteria with a reduction in Verrucomicrobia (**Figure 5F**). Within the taxa firmicutes, there were major perturbations in the order Lactobacillales (**Figure 5G**) and an increase in three lactobacillus species – *johnsonii*, *reuteri-vaginalis*, and *lactis* (**Figure 5H**), which is heavily influenced in aged mice^26^ and associated with reduced colonic inflammation and altered metabolism after SG^36,38^. Further, post SG changes in the duodenum lactobacillus richness is associated with weight loss^37^.

## 4. Discussion

Beyond short-term metabolic benefits, sleeve gastrectomy (SG) has been associated with up to 10 additional years of average lifespan in human cohorts^16^ and exerts broad multi-organ protective effects, including reductions in cardiovascular and renal disease burden, improved pulmonary function, decreased cancer incidence, and enhanced brain health^40^. However, whether these longevity benefits are attributable solely to weight reduction or reflect surgery-specific reprogramming of metabolism — particularly in the context of late-life stressors — remains an unresolved question.

To disentangle surgery-specific metabolic effects from those driven purely by weight loss, our group developed a lean mouse model of SG in which animals are reared and maintained on standard low-fat chow. In this context, we have previously demonstrated that SG improves host glucose metabolism, increases glucose utilization, induces shifts in the adipose tissue transcriptome, and remodels organ-specific immune cell architecture in the absence of any difference in caloric intake, body composition, or weight^14,20^. Building on these observations, we sought to determine whether SG performed in young, lean animals confers lasting protection against metabolic stress encountered in midlife.

Consistent with our prior work and parallel observations following Roux-en-Y gastric bypass (RYGB) in lean mice^41^, SG and sham-operated animals maintained on low-fat chow exhibited identical weight gain, food intake, and near-identical body composition in the postoperative period. Despite this, SG animals displayed markedly altered glucose handling, characterized by accelerated clearance of orally administered glucose^14^. Upon exposure to HFD 60 days after surgery (5 months of age), these differences in glucose handling were accentuated. SG animals demonstrated a consistently attenuated systemic glucose excursion and improved insulin tolerance relative to sham controls, recapitulating findings previously reported following lean RYGB^41^.

In addition to improved glucose metabolism, SG animals demonstrated a unique capacity for metabolic flexibility upon dietary challenge. Indirect calorimetry performed before, during, and after the transition to HFD revealed that SG mice more efficiently switch substrate utilization from carbohydrate to fat oxidation in response to the altered nutrient environment. This enhanced fuel flexibility was accompanied by elevated energy expenditure, resulting in reduced weight gain and adiposity. Notably, the influence of SG on organismal energy balance appears to be diet dependent. We have previously shown that SG animals on a low protein, high carbohydrate diet have a heightened energy expenditure compared to those on standard and high protein diets. However, in the low-protein dietary context, diet, rather than surgery, was the primary driver of energy utilization. In contrast, under conditions of higher dietary protein intake and in the western diet context, SG animals expended more energy than sham counterparts^27^, as observed in the present study. Collectively, these findings support the conclusion that SG uniquely enables metabolic adaptation across diverse dietary exposures.

The augmented fat oxidation observed in SG animals likely carries downstream organ-specific impacts. Consistent with findings in lean RYGB mice^41^, and in animals maintained on lifelong HFD following SG diet^42^, lean SG mice exposed to obesogenic diet in the current study exhibited significantly reduced hepatic lipid accumulation. Although no differences in fibrosis, hepatocellular ballooning, or end-stage steatohepatitis were observed, this is expected given that these histological endpoints are not typically observed in murine models without genetic manipulation, choline-deficient dietary regimens, or extended feeding protocols^31^.

SG-mediated metabolic remodeling dramatically impacts multiple adipose tissue depots. We previously demonstrated that visceral adipose tissue serves as a principal sink for circulating glucose in lean SG mice and that this metabolic reprogramming is accompanied by a shift in the local inflammatory transcriptional signature^14^. Here, we again show that SG modifies the inflammatory state of visceral adipose tissue, reducing pro-inflammatory cytokine expression and attenuating macrophage accumulation upon HFD exposure. Translational relevance is supported by clinical data demonstrating that the suppression of myeloid cell infiltration into adipose tissue following bariatric surgery is surgery-specific and is not recapitulated by lifestyle intervention alone^43^. Thus, our work adds to evidence that bariatric surgery exerts a unique influence over adipose tissue inflammatory phenotype that is independent of weight loss.

The causal mechanisms behind these SG-induced protections remain unclear. The microbiome is an important mediator of metabolism after bariatric surgery. Fecal microbial transplant of stool from humans after bariatric surgery induces weight loss in DIO mice^44^ and antibiotic ablation of the microbiome leads to failure of SG to improve metabolism^45^. In the present study, we identified a large, 3-fold, increase in the total abundance of cecal bacteria following SG. This enrichment in cecal biomass alone likely contributes an outsized influence on the host energy expenditure as baseline measurements of cecal microbiota energy consumption is roughly measured at 8%^46^.

We have previously established that SG effects on glucose metabolism are dependent upon the action of the microbiome^47–49^. In our prior work, cecal microbial composition was characterized by reductions in Verrucomicrobia and Clostridia, and an increase in Lactobacillus^47^, which is remarkably similar to the current study and in independent cohorts^36,37^ despite different diet exposures and a different age at sampling. Among the microbial taxa altered by SG, the order Lactobacillales exhibited the largest change. Lactobacillus species are gram-positive, aerotolerant, anaerobic rods that can be found throughout the gastrointestinal tract and other mucosal surfaces^50^. These bacteria can influence host metabolism through different ways. They produce butyrate and other short chain fatty acids, which are reduced in patients with T2DM. Short chain fatty acids, which are the byproducts of anaerobic fermentation of non-digested carbohydrates, enter circulation and interact with G-protein coupled receptors throughout the body to influence host metabolism^51,52^. Supplementation of lactobacillus species improved fasting glucose^23^ and reduced weight^24^ in humans, improved hyperlipidemia and hepatic steatosis in rats^25^, and suppresses tumor formation in a murine model of non-acholic fatty liver disease-associated hepatocellular carcinoma^53^. The coincident improvement in glucose homeostasis, body weight, and hepatic lipid content in SG animals in the current study may therefore be related, at least in part, to the surgery-associated enrichment of Lactobacillus, though causal attribution will require future investigation.

In the context of SG, Shao and colleagues showed that SG induced a major increase in the duodenal richness of lactobacillus species and through a complex interaction with hypoxia signaling influences host metabolism^37^. Interestingly, Metformin, a first line agent for the treatment of T2DM, also induces enrichment of upper intestinal lactobacillus species which modulates glucose sensing via the sodium-glucose cotransporter-1^54^.

Of particular relevance to aging, a comprehensive longitudinal microbiome analysis in C57BL/6J mice from 2 to 30 months of age found that the microbiome undergoes dramatic restructuring over the lifespan and directly contributes to age-related disease^26^. Strikingly, *Lactobacillus johnsonii*, the Lactobacillus species most enriched in our SG mice at 8 months (**Figure 5H**), was found to decline precipitously between 2 and 9 months of age. This observation raises the intriguing possibility that SG may attenuate age-related microbial decline that would otherwise be expected in middle age and thus, protect metabolic health.

Beyond aging, *L. johnsonii* has been shown to specifically ameliorate diet-induced hypercholesterolemia^55^ and while this study cannot definitively answer whether increased *L*. *johnsonii* in our SG mice is responsible for the improved systemic triglyceride and cholesterol levels, it warrants further study. Additionally, *L. johnsonii* supplementation can improve colonic mucosal integrity and improve colitis phenotypes^56,57^. Given our prior demonstration that SG induces weight-loss-independent shifts in organ-specific immune cell repertoire^20^ and intestinal inflammatory signatures^58^, a mechanistic interaction between the post-SG microbiome, intestinal mucosa, and mucosal immune cells represents a compelling hypothesis to explain the systemic immunological changes observed following surgery, including the reduction in adipose tissue inflammation reported here.

In contrast to the expansion of Lactobacillales, Verrucomicrobia (*Akkermansia*) were reduced following SG in the present study. This is noteworthy because multiple other metabolic interventions like high protein calorie restricted diets^21,22^, time restricted feeding^59^, very low-calorie ketogenic diets^60^, and treatment with the dual GIP/GLP-1 incretin mimic^61^ consistently *Akkermansia* abundance. Further, reductions in *Akkermansia* enrichment across the gastrointestinal tract has been observed in DIO mice, rats, and humans, and *Akkermansia* enrichment correlates with weight loss following fecal microbiota transplantation into obese animals^34,44^ The divergent *Akkermansia* response following SG compared with these other interventions underscores the therapeutic specificity of gut microbiome remodeling and highlights the need for further investigation into the relative contributions of distinct bacterial taxa to the metabolic, aging, and longevity benefits conferred by different therapeutic strategies.

Several limitations of the present study merit consideration. First, all experiments were conducted in male mice. Given well-established sex differences in the efficacy of metabolic therapies, generalizability to female animals and to human populations requires further study. Second, animals were operated on during youth and studied in midlife and whether these protective effects persist into late life, translate to improved longevity and healthspan, or are recapitulated in older animals at the time of surgery remains unknown. That said, clinical evidence indicates that older adults with obesity and metabolic disease do derive meaningful metabolic benefit from bariatric surgery, albeit to a lesser degree than younger patients ^62,63^. Finally, we do not establish a causal relationship between SG, the associated shifts in microbial community structure, and protection against midlife obesity.

## Abbreviations

SG: (Sleeve Gastrectomy),
(T2D): Type 2 Diabetes,
(BMI): body mass index,
(HFD): high fat diet,
(VAT): visceral adipose tissue,
POD: (post-operative Day),
GLP-1: (glucagon like peptide 1),
OGTT: (oral glucose tolerance test),
ITT: (insulin tolerance test),
DIO: (diet induced obesity)

## Credit authorship contribution statement

AM: writing, study execution, and design. RB, TM, DZ: in vitro analysis, writing. SNC, SD: bile acid and microbial analysis. EGS, DAH: Conceptualization, formal analysis, methodology, supervision, writing – review and editing. EGS and DAH are the guarantor of this work and, as such, had full access to all the data in the study and takes responsibility for the integrity of the data and the accuracy of the data analysis.

## Acknowledgements

We would like to thank the Brigham and Women’s Hospital Metabolic Core for their expertise.

## Data Availability

All data is available upon request from the corresponding authors

---

**Supplemental Table 1.**
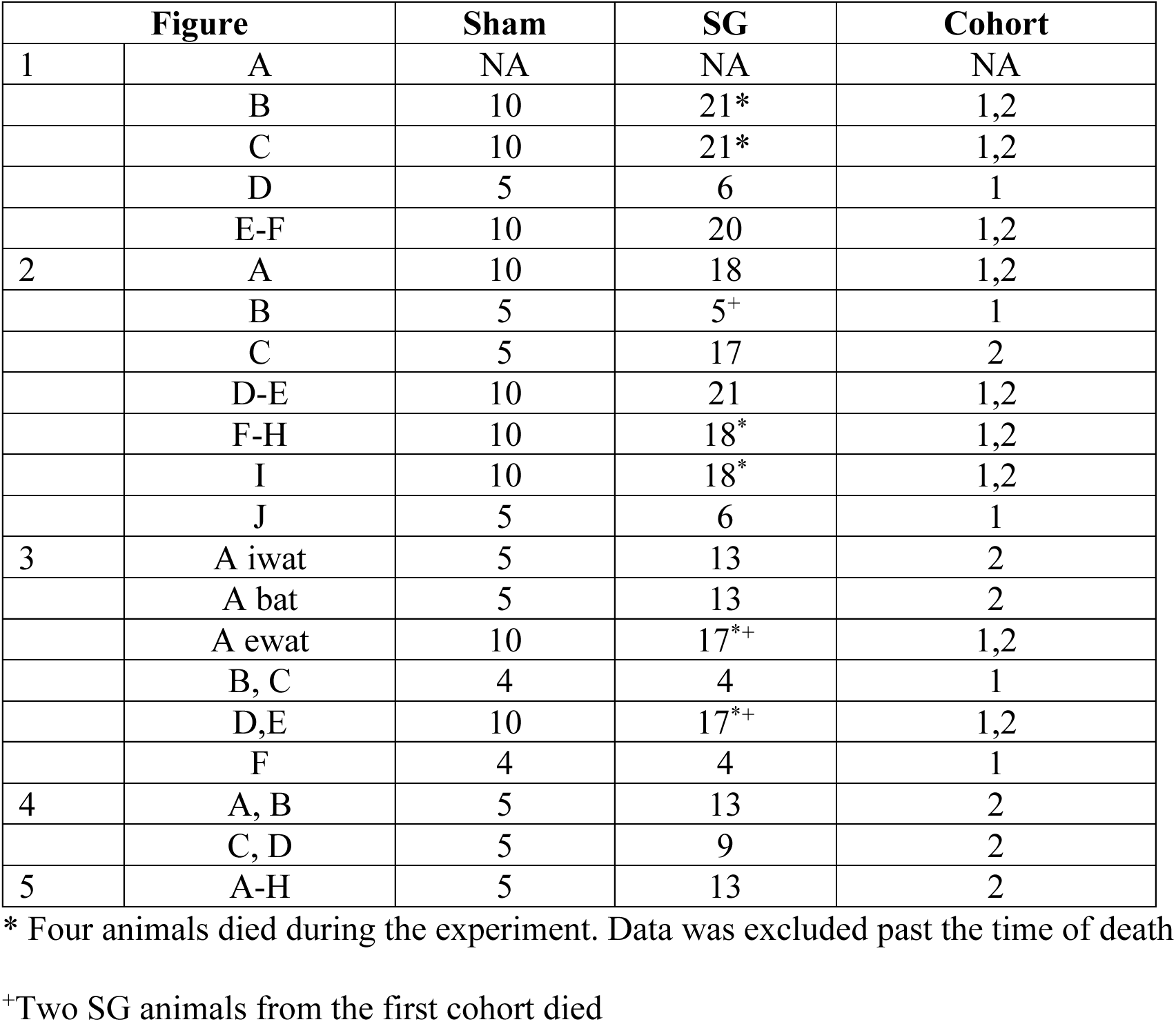
Figure n.

**Supplemental Table 2.** MAFLD scoring.

| NAS Components |  |  |
| --- | --- | --- |
| Item | Score | Extent |
| Steatosis | 0 | <5% |
|  | 1 | 5-33% |
|  | 2 | >33-66% |
|  | 3 | >66% |
| Lobular Inflammation | 0 | No foci |
|  | 1 | <2 foci/200x |
|  | 2 | 2-4 foci/200x |
|  | 3 | >4 foci/200x |
| Hepatocyte ballooning | 0 | None |
|  | 1 | Few balloon cells |
|  | 2 | Many cells/prominent ballooning |
| **Fibrosis Stage** |  |  |
|  | 0 | None |
|  | 1 | Perisinusoidal or peri-portal |
|  | 1A | Mild, zone 3, perisinusoidal |
|  | 1B | Moderate, zone 3, perisinusoidal |
|  | 1C | Portal/periportal |
|  | 2 | Perisinusoidal and portal/periportal |
|  | 3 | Bridging fibrosis |
|  | 4 | cirrhosis |

Total NAS score represents the sum of scores for steatosis, lobular inflammation, and ballooning, and ranges from 0-8. Diagnosis of NASH (or, alternatively, fatty liver not diagnostic of NASH) should be made first, then NAS is used to grade activity. In the reference study, NAS scores of 0-2 occurred in cases largely considered not diagnostic of NASH, scores of 3-4 were evenly divided among those considered not diagnostic, borderline, or positive for NASH. Scores of 5-8 occurred in cases that were largely considered diagnostic of NASH.

**Supplemental Table 3.**
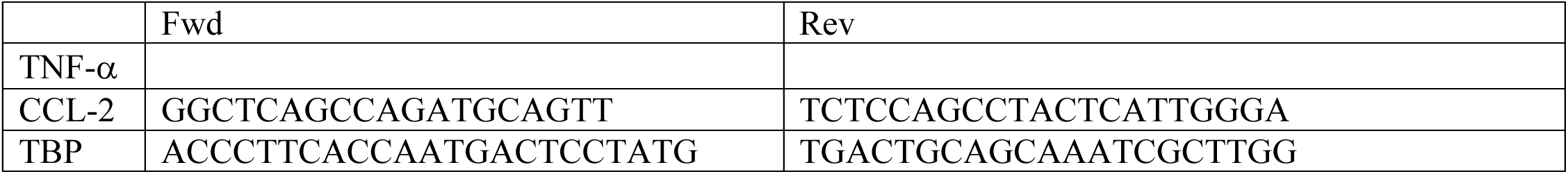
PCR primers.

**Supplemental Table 4.** Analysis of sequential CLAMS runs (p-values reported)

|  | <u>Post-Operative Day 56 to 64 with Diet Crossover</u> |  |  |  |  |  | <u>Post-Operative Day 94-103</u> |  |  |
| --- | --- | --- | --- | --- | --- | --- | --- | --- | --- |
|  | Chow |  |  | HFD |  |  | Post HFD Exposure |  |  |
|  | 24 Hr | Fasting | Feeding | 24 Hr | Fasting | Feeding | 24 Hr | Fasting | Feeding |
| Respiratory Exchange Ratio | 0.480 | 0.107 | 0.610 | 0.056 | **0.048\*** | 0.099 | 0.626 | 0.275 | 0.739 |
| Energy Expenditure (kcal/hr) | 0.647 | 0.962 | 0.398 | 0.713 | 0.247 | 0.952 | **0.012\*** | **0.037\*** | **0.006\*** |
| Hourly Food Consumed (g) | 0.596 | 0.584 | 0.822 | 0.478 | 0.765 | 0.572 | 0.083 | 0.300 | 0.071 |
| Total Food Consumed (g) | 0.757 | 0.805 | 0.725 | 0.136 | 0.158 | 0.305 | 0.326 | 0.314 | 0.340 |
| Locomotor Activity | 0.469 | 0.933 | 0.230 | 0.320 | 0.395 | 0.305 | 0.369 | 0.308 | 0.565 |
| Ambulatory Activity | 0.989 | 0.517 | 0.526 | 0.841 | 0.855 | 0.872 | 0.148 | 0.304 | 0.135 |

